# Length-dependent accumulation of double-stranded RNAs in plastids affects RNA interference efficiency in the Colorado potato beetle

**DOI:** 10.1101/812750

**Authors:** Wanwan He, Wenbo Xu, Letian Xu, Kaiyun Fu, Wenchao Guo, Ralph Bock, Jiang Zhang

## Abstract

Transplastomic potato plants expressing double-stranded RNA (dsRNA) targeted against essential genes of the Colorado potato beetle (CPB) can be lethal to larvae by triggering an RNA interference (RNAi) response. High accumulation levels of dsRNAs in plastids are crucial to confer an efficient RNAi response in the insects. However, whether length and sequence of the dsRNA determine the efficacy of RNAi and/or influence the level of dsRNA accumulation in plastids is not known. Here we compared the RNAi efficacy of different lengths of dsRNA targeted against the CPB *β*–*Actin* gene (*ACT*) by feeding *in vitro*-synthesized dsRNAs to larvae. We show that, while the 60 bp dsRNA induced only a relatively low RNAi response in CPB, dsRNAs of 200 bp and longer caused high mortality and similar larval growth retardation. When the dsRNAs were expressed from the plastid (chloroplast) genome of potato plants, we found that their accumulation levels were correlated with length. dsRNA accumulation levels were positively associated with the observed mortality, suppression of larval growth and suppression of target gene expression. Importantly, transplastomic potato plants expressing the 200 bp dsRNA were better protected from CPB than plants expressing the 297 bp dsRNA, the best-performing line in our previous study. Our results suggest that the length of dsRNAs is an important factor that influences their accumulation levels in plastids and thus determines the strength of the insecticidal RNAi effect. Our findings will aid the design of optimized dsRNA expression constructs for plant protection by plastid-mediated RNAi.

## Introduction

RNA interference (RNAi) is a post-transcriptional gene silencing mechanism trigged by double-stranded RNA (dsRNA) in eukaryotes. dsRNAs are cleaved by endoribonucleases of the Dicer family into 21-23 bp small interfering RNAs (siRNAs) which then are loaded into the RNA-induced silencing complex (RISC) to cleave and degrade cognate mRNAs (Gordon and Waterhouse 2007). Since RNAi was first discovered in the model nematode *Caenorhabditis elegans* (Fire *et al*., 1998), RNAi has been demonstrated to provide a versatile tool in reverse genetics by exploring gene functions through suppression of gene expression (Tomoyasu and Denell 2004; Wheeler *et al*., 2003). In addition, RNAi can be used to modify interactions between organisms, in that dsRNAs produced by one organism can be taken up by another organism and silence genes in that organism, a phenomenon dubbed environmental RNAi or trans-kingdom RNAi. For more than a decade, substantial efforts have been made to develop RNAi-based technologies as a novel approach for insect pest control (Zhang *et al*., 2017). Efficient delivery of insecticidal dsRNAs (targeting essential insect genes) is the key process in RNAi-based pest control. Injection of dsRNA into insects is suitable for laboratory research, but not for practical application towards pest control in the field. Therefore, strategies for oral delivery of dsRNA molecules have been explored in both the laboratory and the field, including the use of nanocarriers, detergents, polymers, engineered bacteria and yeasts, and transgenic plants (He *et al*., 2013; Murphy *et al*., 2016; Zhang *et al*., 2015; Zheng *et al*., 2019; Zhu *et al*., 2011). Since the costs of kilogram amounts of chemically synthesized dsRNA synthesis are very high and large-scale application to plants in the field is burdened with technical problems, expression of dsRNA *in planta* represents the method of choice for the control of herbivorous pest insects. Encouragingly, the first plant-incorporated protectant (PIP) based on RNAi technology (targeting the western corn rootworm *Diabrotica virgifera virgifera*) was approved by the US Environmental Protection Agency (EPA) in 2017 and may enter the market soon (Schiemann *et al*., 2019; Zotti *et al*., 2017).

Although nuclear-transgenic plants expressing dsRNAs targeted against essential insect genes can reduce growth and fitness of pest insects, full protection of the plants and efficient killing of the insects have not been achieved (Baum *et al*., 2007; Mao *et al*., 2007). The reason is that dsRNAs expressed from the nuclear genome are targets of the plant’s own RNAi machinery and efficiently processed into siRNAs by the Dicer endoribonucleases encoded in the plant genome (Gordon and Waterhouse 2007). It has been demonstrated that long dsRNAs are much more effective triggers of gene silencing by environmental RNAi in insects than siRNAs. Feeding studies have revealed that dsRNAs of at least 60 base pairs (bp) in length are necessary for an efficient RNAi response in *Diabrotica virgifera* (Bolognesi *et al*., 2012) and *Tribolium castaneum* (Wang *et al*., 2019). Moreover, since there are no RNA-dependent RNA polymerases in insects, the silencing signals cannot be amplified in insect cells (Price and Gatehouse 2008). Thus, the development of a strong RNAi response in the insect critically depends on the continuous uptake of sufficient amounts of exogenously supplied dsRNA. These limitations explain, at least in part, the limited insect resistance conferred by dsRNA expression in nuclear-transgenic plants.

We recently showed that long dsRNAs can stably accumulate in plastids, a cellular compartment that is derived from formerly free-living cyanobacteria which lack an RNAi pathway (Zhang *et al*., 2015). Transplastomic potato plants expressing dsRNA targeted against the *β*–*Actin* (*ACT*) gene of the Colorado potato beetle (CPB, *Leptinotarsa decemlineata*) were efficiently protected from insect damage and induced a much stronger RNAi response in the beetle than nuclear-transgenic plants expressing a similar dsRNA construct (Zhang *et al*., 2015). This was because long *ACT* dsRNAs (i) accumulate to high levels in plastids owing to the high polyploidy of the plastid genome with up to 10,000 identical copies per cell, and (ii) remained intact due to the absence of an RNAi pathway (and the absence of Dicer-like dsRNA-specific endoribonucleases) from plastids. Although the same promoter (the plastid 16S ribosomal RNA gene promoter, *Prrn*) was used to drive transcription of three different dsRNA constructs in plastids, very different dsRNA accumulation levels were obtained: ∼0.4% of the total cellular RNA for *ACT*, ∼0.05% for *Shrub (SHR)*, and ∼0.1% for *ACT*+*SHR* (fusion of *ACT* and *SHR*). The factors that determine the efficiency of target gene silencing in CPB and the dsRNA accumulation levels in plastids are currently unknown (Table S1).

In the present study, we set out to evaluate the effects of dsRNA length on the efficiency of RNAi as evidenced by the induction of gene silencing and mortality in CPB. We also compared the accumulation levels of dsRNAs of different lengths in transplastomic plants. Our results show that, while all dsRNAs of 200 bp and longer had similar effects on the induction of gene silencing, the dsRNA accumulation levels in plastids were negatively correlated with the length of the dsRNAs. We also established a positive correlation between dsRNA accumulation levels in plastids and RNAi efficacy in CPB.

## Materials and methods

### Insect and plant material and growth conditions

CPB (*L. decemlineata*) larvae and beetles were collected from a potato field in Urumqi (43.82° N, 87.61° E), Xinjiang Uygur Autonomous Region in China. Insects were routinely reared in an insectary at 28 ± 1□ under a 14 h light/ 10 h dark photoperiod and 50-60% relative humidity using leaves from wild-type potato plants as feed. Adult CPBs were allowed to lay eggs which were collected and transferred onto fresh wild-type potato leaves.

Potato (*Solanum tuberosum* cv. Désirée) plants for plastid transformation experiments were grown under aseptic conditions on MS medium supplemented with 30 g/L sucrose (pH adjusted to 5.7) (Murashige and Skoog 1962) and solidified with 0.6% Micro agar (Duchefa). Transplastomic lines were rooted in the same medium and grown to maturity in soil under standard greenhouse conditions.

### In vitro RNA synthesis

dsRNA was synthesized *in vitro* using the T7 RiboMAX™ Express RNAi System (Promega, USA) according to the manufacturer’s instructions. *ACT* fragments of different length were PCR amplified using specific primers containing the T7 promoter sequence (Table S2). The PCR products were purified with a gel extraction kit (Omega, China). For ssRNA synthesis, the *ACT200* fragment was amplified with primer pair act200-F/T7act200Rev (Table S2). *In vitro* transcription reactions were conducted with T7 RNA polymerase (Thermo Scienfic, USA) following the manufacturer’s instructions. The RNA yield was determined with a Nano Photometer (IMPLEN, Germen) and kept at −80□ until further use.

### Insect bioassays

dsRNA feeding bioassays were performed as described previously (Zhu *et al*., 2011). Third-instar CPB larvae were starved for 24 h prior to initiation of feeding assays. Equal amount (16 ng, in 50 μL ddH_2_O) of dsRNAs of different lengths were painted onto fresh potato leaves covering identical areas of 2 × 2 cm, thus adjusting the concentration of dsRNA to 4 ng/cm^2^. ddH_2_O-painted leaves were used as control. Sufficient dsRNA-coated fresh leaves were provided to the insects and exchanged every day for the duration of the bioassays to ensure optimal CPB feeding. Three groups of insects per treatment were investigated, and each group comprised 10 larvae. The body weight of larvae was recorded at day 3, 4 and 5 for all groups. The mortality was recorded daily.

For larval performance assays on transplastomic potato lines, third-instar CPB larvae were allowed to feed on young leaves detached from two-month old transplastomic or wild-type (as control) potato plants. The body weight of the larvae was recorded at the beginning (day 0) of the assay and at day 2, 3 and 4 for all groups. The mortality was recorded daily.

### Quantitative real-time polymerase chain reaction (qRT-PCR)

Total RNA samples were extracted with the RNAiso plus reagent (Takara, Japan). After digestion with gDNA digester (Yeasen, China), cDNA was synthesized using the RevertAid First Strand cDNA Synthesis Kit (Yeasen, China). The reference gene *RP18* was chosen as internal control (Huggett *et al*., 2005; Pfaffl *et al*., 2004). The reaction mixture consisted of 5 µL TB Green Premix Ex Taq II (Tli RNaseH Plus), 1 µL of cDNA and 0.25 µL of forward and reverse primers (Table S2; 10 µM) in a final reaction volume of 10 µL. The qPCR protocol included an initial denaturation step at 95□ for 2 min, followed by 40 cycles of 95□ for 5 s, 60□ for 30 s and 72□ for 30 s. The relative expression of [*-Actin* was calculated by the 2^-ΔΔCT^ method (Livak and Schmittgen 2001; Shi *et al*., 2013). All experiments were repeated in triplicate

### Vector construction

The plastid transformation vectors used in this study were constructed based on the previously described vector pYY12 (Wu *et al*., 2017). The *gfp* cassette in pYY12 was replaced by the dsRNA expression cassette excised from pJZ197 (Zhang *et al*., 2015) by digestion with HincII and NotI, resulting in plasmid pWW1. *ACT* fragments were obtained by PCR amplification with primer pairs ACT700-F/ACT700-R, ACT400-F/ACT400-R and ACT200-F/ACT200-R, respectively (Table S2), using a clone containing the full-length *β-Actin* cDNA as template. The resulting PCR products were digested with SbfI and SacI, and cloned into the similarly cut vector pWW1, generating plastid transformation vectors pWW3, pWW4 and pWW7, respectively.

### Potato plastid transformation

Potato plastid transformation was performed as described previously (Zhang *et al*., 2017). Briefly, young leaves from aseptically grown potato plant were bombarded with DNA-coated 0.6 µm gold particles using a PDS-1000/He Biolistic Particle Delivery System (Bio-Rad, Hercules, CA, USA). Plasmid DNA for plastid transformation was prepared by using the Nucleobond Xtra Plasmid Midi Kit (Macherey-Nagel, Germany). Spontaneous antibiotic-resistant mutants were eliminated by screening on shoot regeneration medium supplemented with both spectinomycin and streptomycin (Bock 2001). Transplastomic events were confirmed by Southern blot analyses and subsequently purified to homoplasmy.

### Isolation of nucleic acids and gel blot analysis

Total DNA from wild-type and transplastomic plants was extracted by a cetyltrimethylammonium bromide (CTAB)-based method (2% CTAB, 1.4 mM NaCl, 0.1 mM Tris–HCl pH8.0, 20 mM EDTA pH8.0) (Doyle and Doyle 1990). Total RNA was isolated using the RNAiso plus reagent following the manufacturer’s instruction (Takara, Japan). For Southern blot analyses, samples of 5 µg total DNA were digested with the restriction enzymes AgeI and MluI for 12-16 h, separated by electrophoresis in 1% agarose gels, and transferred onto Hybond nylon membranes (GE Healthcare) by capillary blotting. A PCR product covering a portion of the *psbZ* coding region (You *et al*., 2019) was used as a hybridization probe.

For RNA gel blot analyses, RNA samples were denatured, separated in formaldehyde-containing 1% agarose gels and blotted onto Hybond nylon membranes (GE Healthcare). A PCR product generated by amplification of *ACT* cDNA with primer pair act-N-F/act-N-R served as probe to determine dsRNA amounts (Table S2). The probe was labeled with the DIG-High Prime DNA Labeling and Detection Starter Kit II following the manufacturer’s instructions (Roche, USA).

### Statistical analysis

Survival curves were analyzed using the Kaplan-Meier method. The log-rank test was used to evaluate the significance of differences between two groups. qRT-PCR data were used to calculated the relative expression of *β-Actin* by the 2^-ΔΔCT^ method. Data of qRT-PCR and body weight were analyzed with one-way ANOVA coupled with Bonferroni (equal variances) or Dunnett’s T3 (unequal variances) correction multiple comparison test. A value of P<0.05 was considered significantly different. Data were statistically analyzed by SPSS version 19.0. Figures were drawn by GraphPad Prism 7 and SnapGene 3.2.1.

## Results

### Effect of dsRNAs of different lengths on RNAi efficacy in CPB larvae

To avoid potential effects from different GC contents of the selected dsRNA target regions, we cloned *β-Actin* fragments (*ACT*s) of different lengths (60, 200, 297, 400 and 700 bp) but similar GC content (∼52.2%). The cloned gene fragments were then used as templates for dsRNA synthesis by *in vitro* transcription (Fig. S1). To assess the effects of ds*ACT* length on RNAi efficacy in CPB larvae, ds*ACT*-painted leaves with defined amounts of dsRNA (4 ng/cm^2^) were fed to third-instar larvae. Compared to the control (fed with H_2_O-painted leaves), feeding of larvae with ds*ACT* of lengths ranging from 200 bp to 700 bp resulted in significantly increased mortality, inhibited growth and suppressed *β-Actin* expression (Fig. 1). No significant differences in killing efficiency, weight gain of survivors and level of gene silencing were detected between ds*ACT*s of lengths between 200 and 700 bp. By contrast, ds*ACT60* was much less effective than ds*ACT*s of 200 bp and greater length. Compared to the control, only slightly retarded larval growth was observed (Fig. 1C), but no significantly increased mortality and no significant target gene silencing. In summary, the RNAi effects obtained with ds*ACT*s of different length were shown to be 200 bp ≈ 297 bp ≈ 400 bp ≈ 700 bp > 60 bp.

**Fig. 1.**
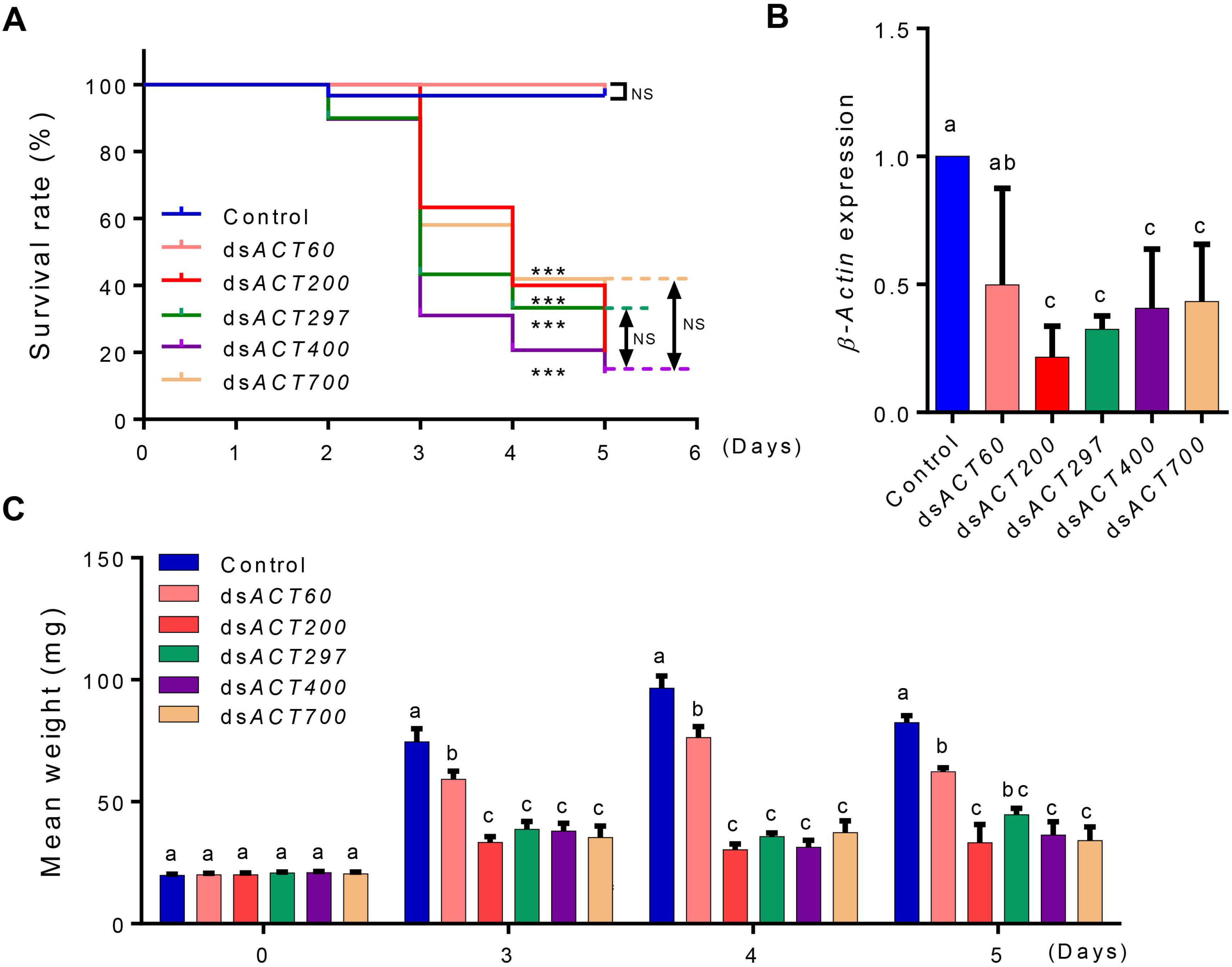
RNAi effects on CPB larvae fed with different length of i*n vitro* synthesized dsRNAs. (A) Kaplan-Meier survival curves of third-instar CPB larvae fed with potato leaves that had been painted with identical amounts (4 ng/cm^2^) of ds*ACTs* of different lengths. The log-rank test was used to assess the significance of differences between two survival curves. \*\*\**P* < 0.001; NS, not significant. (B) Relative expression levels of *β-Actin* in the CPB larvae in (A) at day 5. Gene expression levels were set as one in CPB larvae fed with H_2_O-painted control leaves. Data are means ± SD (n = 3). (C) Mean weight of surviving CPB larvae at the indicated days of feeding. Data are means ± SE (n = 6). The letters above each bar in (B) and (C) indicate the significance of differences as determined by one-way ANOVA in SPSS (Bonferroni’s test).

### Generation and molecular characterization of transplastomic potato plants expressing dsACTs

Having seen very similar RNAi effects of ds*ACT*s above a length of 200 bp on CPB, we next constructed potato plastid transformation vectors for expression of ds*ACT200* (vector pWW7), ds*ACT400* (vector pWW4) and ds*ACT700* (vector pWW3) (Fig. 2A). The constructs were introduced into the potato plastid genome by particle gun-mediated (biolistic) chloroplast transformation. The expression of ds*ACT*s in plastids is driven by two convergent copies of the tobacco plastid ribosomal RNA operon promoter (*Prrn*). The plastid-specific selectable marker gene *aadA* (conferring spectinomycin resistance) is driven by the chloroplast *psbA* promoter and the 3’ untranslated region of the *rbcL* gene from the unicellular green alga *Chlamydomonas reinhardtii*. Selection for spectinomycin resistance, several independent transplastomic potato lines were obtained for each construct, and two independent transplastomic lines per construct were selected for further characterization. Transgene integration into the plastid genome by homologous recombination was confirmed by Southern blot analysis. The homoplasmic status of the transplastomic potato lines (i.e., the absence of residual wild-type copies of the highly polyploid plastid genome) was evidenced by the absence of a hybridization signal for the 2.13 kb fragment diagnostic of the wild-type plants (St-wt) and exclusive presence of the 2.51 kb fragment expected in transplastomic lines (Fig. 2B).

**Fig. 2.**
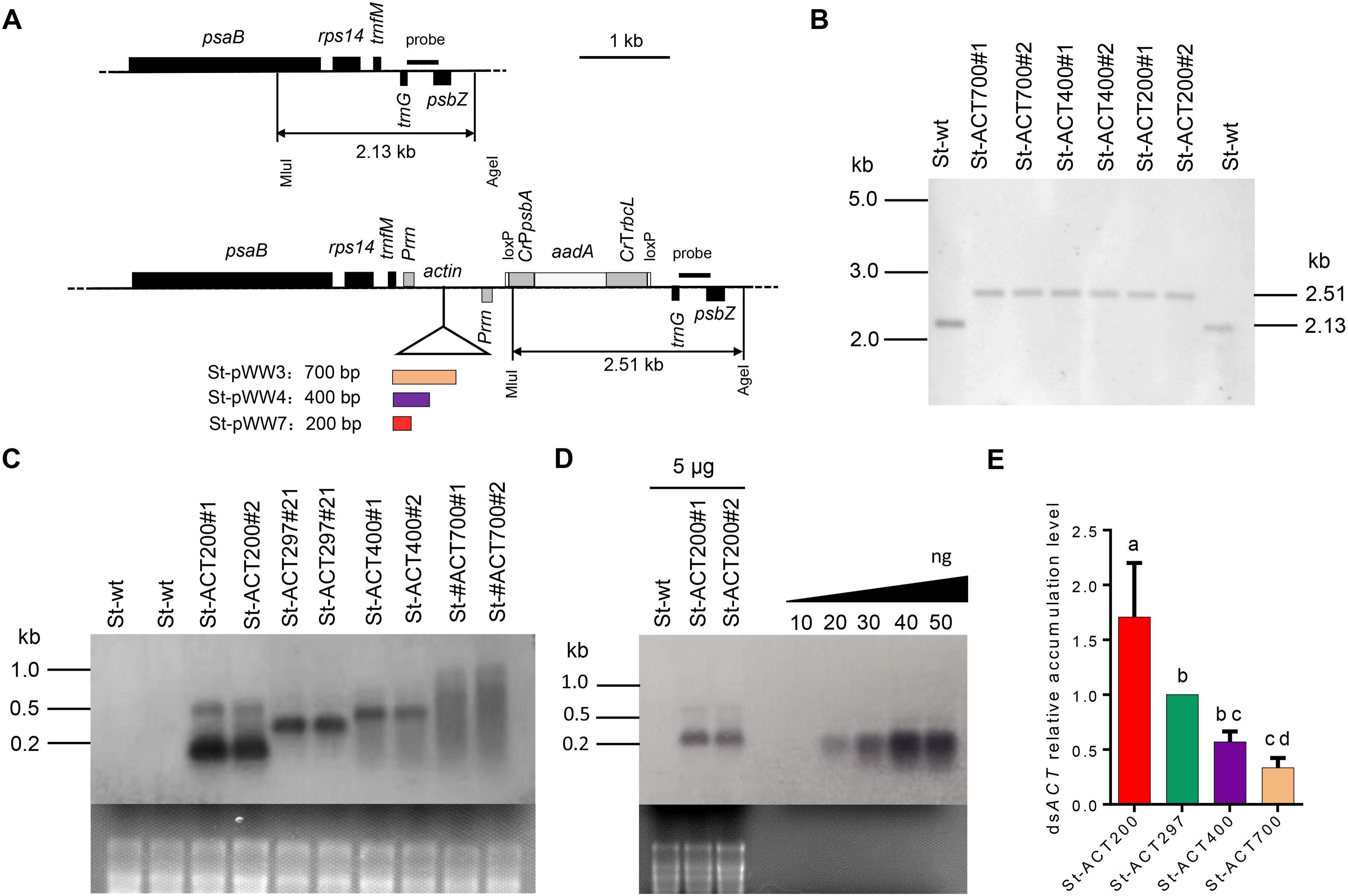
Generation of transplastomic potato plants and analysis of ds*ACT* accumulation in plastids. (A) Physical maps of the targeting region in the plastid genome of wild-type potato and map of the transformation vectors for ds*ACT* expression from the plastid genome. The dsRNA expression cassettes contain two convergent *Prrn* promoters for dsRNA production. The selectable marker gene *aadA* is driven by the *psbA* promoter and the 3′ untranslated region of the *rbcL* gene from *Chlamydomonas reinhardtii*. Genes above the line are transcribe from left to right, genes below the line are transcribed in the opposite direction. The MluI and AgeI restriction sites used for RFLP analysis of transplastomic lines are indicated, and the sizes of the restriction fragments detected in Southern blot analyses are given. The location of the hybridization probe is shown as a black bar. (B) Southern blot confirming homoplasmy of the transplastomic potato lines. Absence of the 2.13 kb hybridization signal for the wild-type genome indicates homoplasmy of transplastomic lines. (C) Analysis of ds*ACT* accumulation by northern blotting. 5 μg of total cellular RNA were loaded in each lane. The ethidium bromide-stained gel prior to blotting is shown below the blot. (D) Quantification of dsRNA accumulation levels in St-ACT200 transplastomic potato lines by northern blot analyses. Samples of 5 μg total cellular RNA were loaded for the wild-type (St-wt) and two independent St-ACT200 lines. A dilution series of *in vitro* synthesized ssRNA was loaded for comparison. (E) Relative quantification of the expression levels of ds*ACT*s of different lengths in plastids. The signal intensities of the hybridizing bands of the expected size in ds*ACT* northern blots (cf. panel c) were quantified using the ImageJ software. Values are normalized to St-ACT297 (set to 1.0). Data represent the means of four independently performed northern blot experiments.

### Determination of dsRNA accumulation levels in plastids

Having obtained the transplastomic potato plants, we next examined the accumulation levels of the ds*ACT*s of different lengths in plastids by northern blot analysis. For comparison, we included the previously generated transplastomic potato line St-ptDP-ACT21 (expressing dsA*CT297*; referred to as St-ACT297) (Zhang *et al*., 2015). The data revealed that the accumulation levels of ds*ACT*s in the different transplastomic lines were St-ACT700 < St-ACT400 < St-ACT297 < St-ACT200. These findings indicate a length-dependent mode of dsRNA accumulation in plastids in that the accumulation levels decrease with the length of the dsRNA (Fig. 2C). When compared to a dilution series of *in vitro* synthesized RNA, ds*ACT* accumulation levels in St-ACT200 line were estimated to be ∼0.6% of the total cellular RNA (Fig. 2D), which is ∼1.5 times as high as in St-ACT297 (∼0.4% of the total cellular RNA) (Fig. 2E).

We also noted evidence of substantial dsRNA degradation in the transplastomic lines expressing longer dsACT fragments (Fig. 2C), suggesting that the lower levels of ds*ACT* accumulation in these plants are caused by RNA instability. This interpretation is also consistent with all constructs being driven by identical promoters, thus likely resulting in identical rates of transcription.

### RNAi effects of transplastomic potato plants on CPB

We next evaluated whether there were differences in the level of insect resistance conferred by the transplastomic potato lines expressing different lengths of ds*ACT*. Third-instar CPB larvae were chosen for the insecticidal bioassays, because they display relatively high survivorship compared to first- or second-instar CPB larvae when exposed to lethal dsRNAs (Guo *et al*., 2015).

Our bioassays revealed that, compared to the wild-type control (St-wt), feeding of CPB with leaves from St-ACT200, St-ACT297 and St-ACT400 lines resulted in high mortality of the larvae. The CPB larvae fed with these lines showed similarly strong reductions in weight gain (Fig. 3C). Interestingly, starting from day 2, the larvae almost completely ceased feeding on the St-ACT200, St-ACT297 and St-ACT400 plants (Fig. S2). By contrast, no significant mortality was observed in CPB larvae fed on the St-ACT700 line that showed the lowest ds*ACT* accumulation levels (Fig. 2C,3A). Although resulting in reduced larval weight gain and detectable induction of gene silencing in CPB larvae compared to the control (feeding on wild-type plants), the St-ACT700 line causes a much weaker RNAi effect on the CPB larvae than the other St-ACT lines (Fig. 3B,C). The RNAi effects on CPB larvae fed with St-ACT transplastomic plants were St-ACT700 < St-ACT400 < St-ACT297 < St-ACT200, and thus are strikingly correlated with the ds*ACT*s accumulation levels in plastids (Fig. 2C). The St-ACT200 plants caused even higher rates of killing of CPB larvae than the St-ACT297 plants, the best-performing transplastomic potato plants reported previously (Zhang *et al*., 2015) (Figure 3a, S2).

**Fig. 3.**
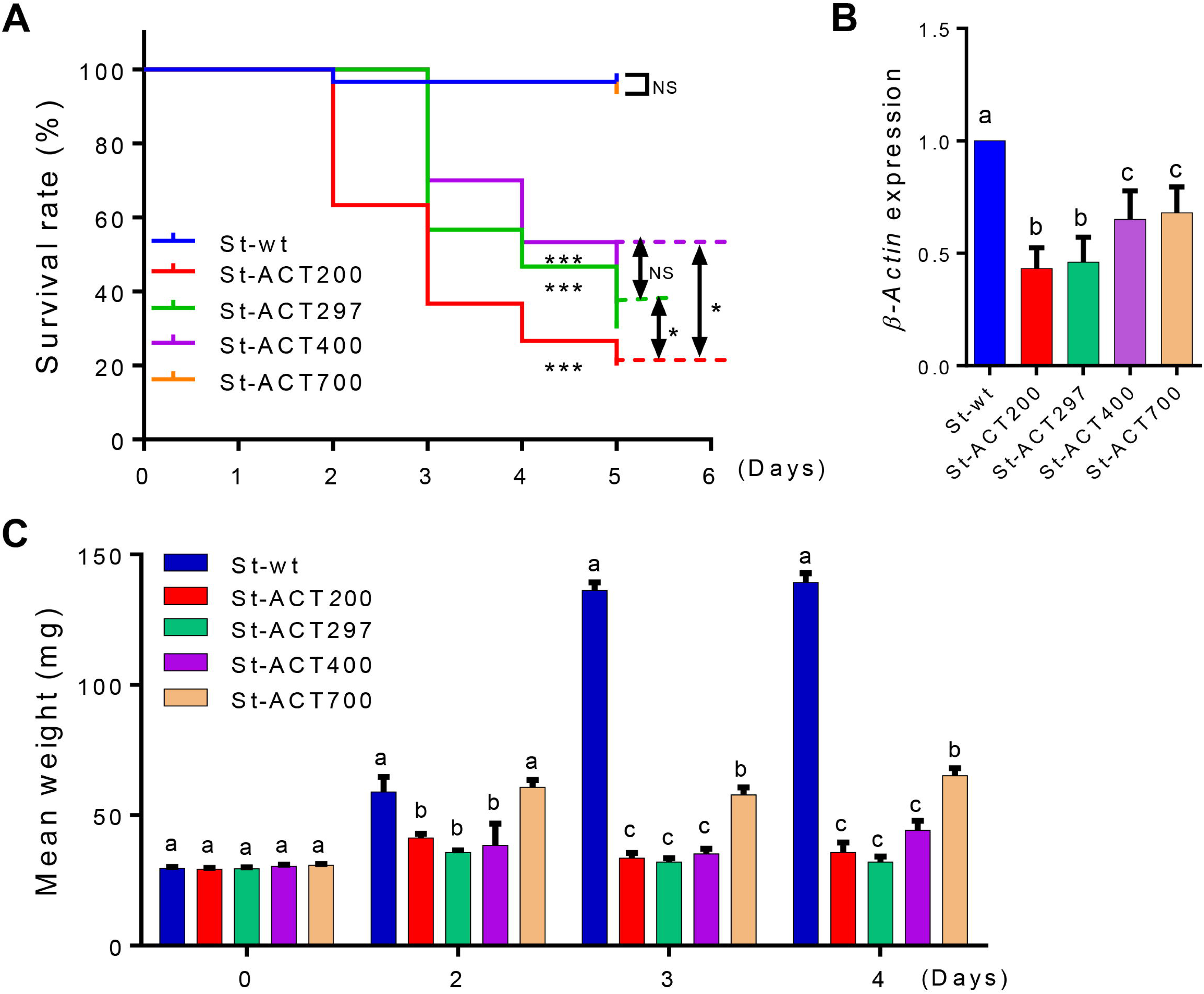
Feeding assays of CPB larvae on transplastomic potato plants. (A) Kaplan-Meier survival curves of third-instar CPB larvae upon feeding on detached leaves of St-ACT lines. The log-rank test was used to assess the significance of differences between two survival curves. \*\*\**P* < 0.001, \**P* < 0.05; NS, not significant. (B) Relative expression levels of *β-Actin* in CPB larvae at day 3 of the assay. Gene expression levels were set as one in CPB larvae fed with wild-type (St-wt) potato plants. Data are presented as means ± SD. n = 5. (C) Mean weight of CPB larvae at the indicated days of feeding. Data are presented as means ± SE. n = 30. The letters above each bar in (B) and (C) indicate the significance of differences as determined by one-way ANOVA in SPSS (Bonferroni’s test, *P* < 0.05).

## Discussion

RNAi has emerged as a promising strategy for insect pest control. However, the factors influencing the efficiency of environmental RNAi in insects are not fully understood. The length of the dsRNA trigger has been demonstrated to be an important factor that affects RNAi efficacy in *T. castaneum* (Miller *et al*., 2012; Wang *et al*., 2019) and *D. virgifera* (Bolognesi *et al*., 2012). It was reported that dsRNAs of at least 60 bp in length are required for efficient RNAi. 21 nt short interfering RNAs (siRNAs), the products of dsRNA processing by Dicer, had little effect on *D. virgifera*. Similarly, dsRNAs of 69 bp, but not of 30 bp, were reported to induce the RNAi responses in *T. castaneum* (Miller *et al*., 2012). A number of factors, including the cellular uptake efficiency of dsRNAs and dsRNA degradation in the midgut of insects, have been implicated in the length-dependent effects of dsRNA. Based on the established threshold length of ∼60 bp, we tested dsRNAs of lengths equal to or greater than 60 bp that target the CPB *β-Actin* gene (ds*ACT*s). Consistent with previous studies, we found that ds*ACT*s of 200 bp and longer are much more potent than dsRNA of 60 bp. Remarkably, no significant increase in RNAi efficiency was observed with dsRNAs of lengths greater than 200 bp (Fig. 1). The finding that expression of very long dsRNAs does not improve the insecticidal activity is important and should be considered in future efforts to design optimized RNAi constructs for pest control.

We previously showed that plastids can be engineered to produce dsRNA to control CPB. We also reported that different dsRNAs can accumulate to different levels in chloroplasts (Zhang *et al*., 2015). Plastid genes are transcribed by two different RNA polymerases in angiosperms: a phage-type nucleus-encoded RNA polymerase (NEP), and a eubacterial-type plastid-encoded RNA polymerase (PEP). Two RNA polymerases can jointly or individually transcribe different plastid genes (Borner *et al*., 2015; Hajdukiewicz *et al*., 1997). The *Prrn* promoter used to drive dsRNA expression in this study is mainly transcribed by PEP (Suzuki *et al*., 2003). Our results indicate that dsRNA accumulation levels decrease with increasing dsRNA length (Fig. 2C). As the same expression signals (*Prrn*) were used to produce ds*ACT*s in all of our transplastomic lines, it seems reasonable to assume that the different (length-dependent) accumulation levels of ds*ACT*s are not due to different rates of transcription, but rather have a post-transcriptional cause. Possible reasons for the length-dependent accumulation of dsRNAs could be related to the efficiency of dsRNA formation (by annealing of the two complementary single strands) and/or decreasing dsRNA stability with length. So far, no dsRNA processing or degrading enzymes have been found in plastids (Stern *et al*., 2010). We, therefore, suspect that the accumulation levels of ds*ACT*s in plastids is more likely to be determined by the efficiency of the dsRNA annealing process. dsRNA formation from single strands is conceivably negatively correlated with length, in that long single-stranded RNAs have a higher propensity to form secondary structures than short ones, which may impede perfect annealing to dsRNAs. Imperfectly annealed single strands may form loops and bulges that then are attacked by endoribonucleases present in plastids.

However, our data reported here do not fully explain the previous observation that three types of dsRNAs (ds*ACT*, ds*SHR* and ds*ACT*+*SHR*) accumulated to very different levels in plastids (Zhang *et al*., 2015) (Table S1). For example, ds*SHR* (220 bp in length) was shorter than ds*ACT* (297 bp in length), while the accumulation level of ds*SHR* in plastids was at least eight-fold lower than that of ds*ACT*. This suggests that dsRNA length is not the only factor determining the accumulation levels of dsRNAs in plastid. Whether other factors like primary sequence and GC content of dsRNAs influence dsRNAs accumulation in plastids remains to be investigated. Conceivably, these factors could also affect the efficiency of strand annealing and, in this way, act similarly as discussed above for RNA length.

Since insects have no RNA-dependent RNA polymerase, the RNAi efficiency in insects is believed to be largely dependent on the amount of environmental dsRNAs that can be delivered to and taken up by insect cells (Price and Gatehouse 2008). This assumption is consistent with our results that the RNAi effects of St-ACT plants on CPB were strongly positively correlated with dsA*CT* accumulation levels in plastids (Fig. 2C,3). ds*ACT200* was found to accumulate to the highest level (∼0.6% of total cellular RNA in St-ACT200 lines; Fig. 2C,D) and resulted in of the strongest RNAi effects in CPB larvae (Zhang *et al*., 2015). Thus, shortening the length of dsRNAs expressed in transplastomic plants may be a useful strategy to increase dsRNA accumulation levels in plastids and, in this way, enhance their insecticidal effect.

Taken together, our data revealed that dsRNA length is an important factor that determined dsRNA accumulation levels in plastids, and influences the efficacy of the RNAi response triggered by transplastomic plants in pest insects. This study will facilitate the design of optimized expression constructs to maximize dsRNA accumulation levels in plastids and increase the efficacy of plastid-mediated RNAi for pest control.

## Acknowledgements

This work was supported by the National Natural Science Foundation of China (31572071). We also thanks Dr. Peng Han (Xinjiang Institute of Ecology and Geography, CAS) for assistance with potato plants cultivation.

## Author contributions

JZ conceived the project; WH and WX performed the research; LX and JZ analyzed the data. KF and WG support the research and provided the CPB resources. RB and JZ wrote the article with contributions of all other authors.

